# Imaging beyond the super-resolution limits using ultrastructure expansion microscopy (UltraExM)

**DOI:** 10.1101/308270

**Authors:** D. Gambarotto, F. U. Zwettler, M. Cernohorska, D. Fortun, S. Borgers, J. Heine, J. G. Schloetel, M. Reuss, M. Unser, E. S. Boyden, M. Sauer, V. Hamel, P. Guichard

## Abstract

For decades, electron microscopy (EM) was the only method able to reveal the ultrastructure of cellular organelles and molecular complexes because of the diffraction limit of optical microscopy. In recent past, the emergence of superresolution fluorescence microscopy enabled the visualization of cellular structures with so far unmatched spatial resolution approaching virtually molecular dimensions. Despite these technological advances, currently super-resolution microscopy does not permit the same resolution level as provided by electron microscopy, impeding the attribution of a protein to an ultrastructural element. Here, we report a novel method of near-native expansion microscopy (UltraExM), enabling the visualization of preserved ultrastructures of macromolecular assemblies with subdiffraction-resolution by standard optical microscopy. UltraExM revealed for the first time the ultrastructural localization of tubulin glutamylation in centrioles. Combined with super-resolution microscopy, UltraExM unveiled the centriolar chirality, an ultrastructural signature, which was only visualizable by electron microscopy.

## INTRODUCTION

Cells are constituted of organelles, tiny elements that carry specific roles for the correct functioning of cells. At the nanometer scale, organelles are large macromolecular assemblies that display specific structures corresponding to their cellular function. Due to their dimensions and the diffraction limit of optical microscopy, electron microscopy was for decades the only method capable of visualizing structural details of the inner life of cells^1^. In the last decade, superresolution microscopy has evolved as a very powerful method for subdiffraction-resolution fluorescence imaging of cells and structural investigations of cellular organelles^2,3^. Super-resolution microscopy methods, such as stimulated emission depletion (STED) microscopy^4^, structured illumination microscopy (SIM)^5^, photoactivated localization microscopy (PALM)^6^ and *direct* stochastic optical reconstruction microscopy (*d*STORM)^7^, now can provide spatial resolution that is well below the diffraction limit of light microscopy, enabling invaluable insights into the spatial organization of proteins in biological samples at the nanometer scale. Despite these advances, the practical resolution of super-resolution microscopy remains limited to ~30-50 nm using standard immunolabeling methods with primary and secondary antibodies with a size of ~ 10-20 nm^8^. The use of camelid antibodies, termed nanobodies or chemical tags such as SNAP-tags, reduces this linkage error but labeling efficiency and specificity still cause problems^9^. In addition, the extractable structural information from super-resolution microscopy data is not only determined by the size of the label and the optical resolution of the instrument but, equally, by the labeling density as dictated by the Nyquist-Shannon sampling theorem^10^. Finally, as super-resolution microscopy is approaching electron microscopy resolution, the method is facing similar technical limitations such as nanometer deformation of the sample due to chemical fixation and permeabilization artifacts^11–13^. Thus, visualization of ultrastructural details of macromolecular assemblies by superresolution microscopy remains challenging^14^.

Recently an innovative method emerged, named expansion microscopy (ExM), which physically expands the fluorescently immunolabeled sample before imaging and thus enables super-resolution imaging by standard fluorescence microscopy^15^ (Fig. 1a). ExM degrades native proteins by strong proteinase treatment to achieve free volumetric expansion of the embedded sample. Hence, only fluorescent tags covalently linked to the dense hydrophilic (acrylic) polymer network are physically separated during expansion still maintaining the overall three-dimensional (3D) imprint of the native biological structures. Upon polymer expansion, the sample expands isotropically ~4.5-fold, enabling ~70 nm lateral resolution using diffraction-limited microscopes^15–18^. While this innovative method in combination with superresolution microscopy might pave the way towards fluorescence imaging with so far unmatched spatial resolution, it also faces technical limitations. One of them is fluorophore loss during the polymerization and protease digestion step^16^. In combination with the lower fluorophore density due to physical expansion, it substantially reduces the overall fluorescence intensity of the sample and consequently also the achievable structural resolution (Fig. 1a, **ExM**). Finally, the distance of the fluorescent probes, usually linked to an IgG antibody, relative to the epitope to be revealed, again impedes high-resolution imaging.

**Figure 1.**
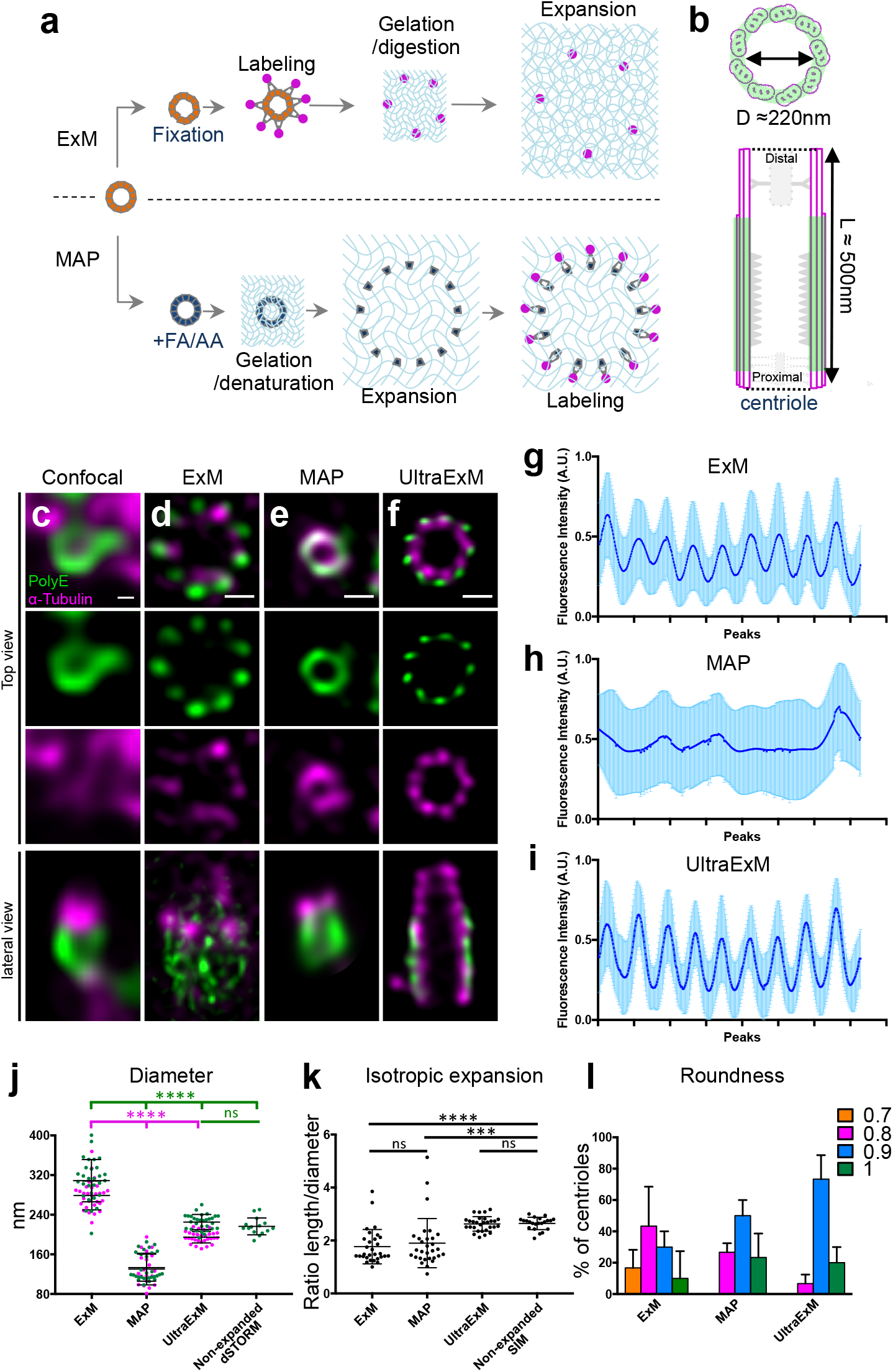
Centriole expansion using UltraExM. (a) Schematic illustration of two methods of expansion microscopy, ExM and MAP. **(b)** Schematic representation of a centriole seen either in top view (top) or lateral view (bottom), **(c-f)** Non-expanded **(c)** and expanded **(d-f)** isolated centrioles stained with PolyE (green, Alexa488) and α-tubulin (magenta, Alexa568) imaged by confocal microscopy followed by HyVolution. Centrioles were expanded using ExM (d), MAP (e) or UltraExM (f). Scale bar in c: 100nm and d-f: 450 nm, (g-i) Plot profile of the polar transform showing the 9-fold symmetry for ExM (g), MAP (h) and UltraExM (i). (j) Diameter of the centrioles in the different conditions. Green and magenta dots represent PolyE and α-tubulin diameters, respectively. PolyE: 308 nm ± 8 nm, 133 nm ± 5 nm, 225 nm ± 3 nm and 216 nm ± 4 for ExM, MAP, UltraExM and Nonexpanded *d*STORM respectability. α-tubulin: 279 nm ± 5, 130 nm ± 7 nm and 195 ±2, for ExM, MAP and UltraExM respectability. (k) Isotropic expansion measured as the ratio between the centriolar length and diameter: ExM=1.8, MAP=1.9, UltraExM=2.6, Non-expanded SIM=2.6. (l) Roundness, shape of the centriole for the three expansion methods. ns=non significant, ***(P=0.0002), ****(P<0.0001). Note that for all the quantifications provided in this figure, we included data from UltraExM performed with 0.7%FA + 0.15% and 0.7%FA + 1% AA.

Alternative protocols of ExM such as Protein Retention ExM (ProExM) and Magnified Analysis of the Proteome (MAP) have been developed that cross-link proteins themselves in the polymer matrix. Replacing protein digestion by heat and chemical induced denaturation allows post-expansion immunostaining of chemically embedded proteins, which still preserve epitope sites for antibody recognition (Fig. 1a, **MAP**). By using retention of intact proteins, followed by post-expansion labeling, these methods benefit from the advantage that the fluorescent labels are then proportionally smaller compared to the size of the expanded sample. This effectively decreases the relative distance of the fluorescent label to the epitope. While ProExM still needs proteolysis for reliable expansion^18^, MAP keeps an intact proteome^17^. MAP is mainly based on preventing the intra- and interprotein crosslinking by incubating the specimen with high paraformaldehyde and acrylamide monomers (PFA/AA) prior to gelation. Acrylamide-bound proteins are then crosslinked to the swellable polymer during polymerization and subjected to denaturation and expansion prior to labeling (Fig. 1a). While MAP should in principle reduce the linkage error, it remains unclear whether it preserves the molecular architecture of organelles or macromolecular assemblies.

Here, we report the development of ultrastructure ExM (UltraExM), a novel expansion microscopy technique that preserves the molecular architecture of multiprotein complexes enabling super-resolution imaging of ultrastructural details by optical microscopy. Importantly, UltraExM coupled to STED microscopy and applied on purified centrioles, unveils for the first time two major hallmarks of centrioles, the 9-fold symmetry of the microtubule triplets as well as their chirality, two ultrastructural features that have so far only resolvable by electron microscopy.

## RESULTS

### Preserving centriole architecture using UltraExM

We first set out to characterize the macromolecular expansion performance using established ExM and MAP protocols^16,17^. To validate the performance of the different protocols in macromolecular expansion, we used isolated *Chlamydomonas* centrioles as reference structure with known precise dimensions^19^. Centrioles are evolutionary conserved organelles important for fundamental cell biological processes such as centrosome and cilia assembly. They are characterized by a nine-fold microtubule triplet-based radial symmetry, forming a polarized cylinder with typical ~ 500 nm long and ~ 250 nm wide dimensions that are close to the diffraction limit of optical microscopy and, as such represents a challenge to image its structural elements (Fig. 1b). Isolated centrioles from the green algae *Chlamydomonas reinhardtii* were immunolabeled for α-tubulin to visualize the centriolar microtubule wall and for polyglutamylated tubulin (PolyE), a tubulin modification present only on the central region of the centriole^20^. While the cylindrical nature of the centriole is visible with the PolyE signal in confocal fluorescence microscopy, it is impossible to reveal the canonical 9-fold symmetry of the microtubule triplets (Fig. 1c). Moreover, we also noticed an antibody competition when co-staining for both α-tubulin and PolyE, both antibodies recognizing epitopes on the C-teminus moiety of tubulin, impeding proper visualization of tubulin in centrioles (Fig. 1c).

Next, we expanded centrioles using both the ExM and MAP protocol and imaged the samples by confocal fluorescence microscopy (Fig. 1d-e). The gels expanded ~4.2-fold using ExM and ~3.5-fold using MAP. Surprisingly, we noticed that the diameter of the centriole in ExM was markedly larger than expected from the determined expansion factor. Indeed, the PolyE signal depicts an average diameter of 308 nm ± 8 nm after expansion compared to the diameter of 216 nm ± 4 nm determined from non-expanded centrioles imaged by *d*STORM using the same primary antibodies (Fig. 1j, **Supplementary Fig. 1a**). This corresponds to a 1.4x enlargement of the expected centriole diameter after expansion suggesting some defects in the macromolecular expansion of centrioles. Moreover, the tubulin signal appears inhomogeneous (Fig. 1d, lateral view and **Supplementary Fig. 2a**). We hypothesized that this could be due to epitope masking of anti-PolyE antibodies as already observed by confocal fluorescence microscopy without expansion (Fig. 1c). Indeed, when centrioles were stained with only α-tubulin, the microtubule triplets were clearly visible (Supplementary Fig. 2b). Moreover, we noticed that the 9-fold symmetry of centrioles can be visualized, albeit not perfectly in the ExM treated centrioles (Fig. 1d, 1g and **Supplementary Fig. 1e**). In contrast, we observed that the MAP treated centrioles appeared 1.6x smaller with an average diameter of 133 nm ± 5 nm (Fig. 1e and Fig. 1j), suggesting as well a defect in the macromolecular expansion of centrioles. As a consequence, the 9-fold symmetry of the PolyE was not apparent (Fig. 1e, 1h). However, in comparison to the confocal and the ExM conditions, we observed a reduced antibody competition issue (Fig. 1e).

**Figure 2.**
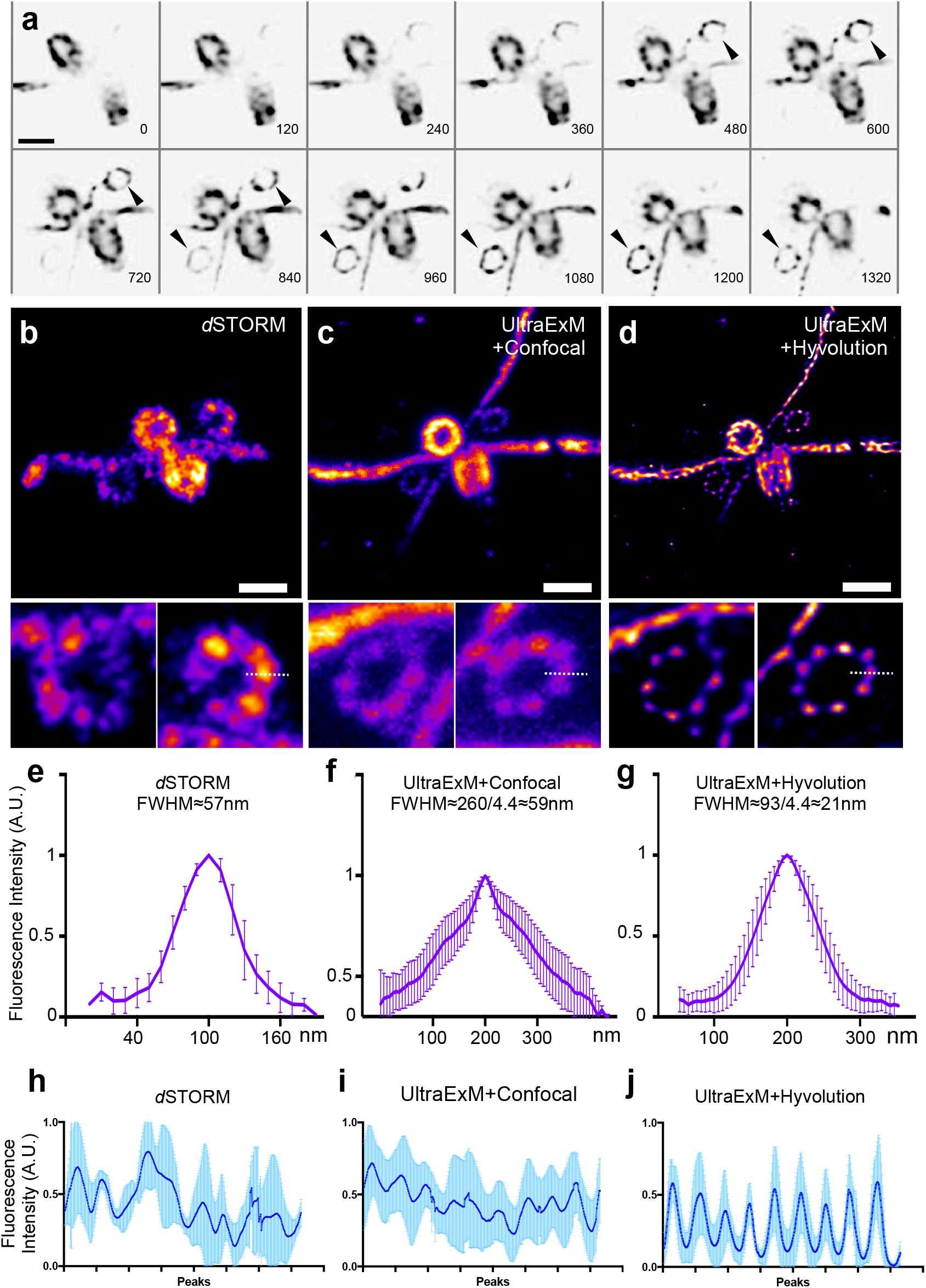
UltraExM reaches the *d*STORM precision limits. **(a)** Montage of a Z-stack through *Chlamydomonas reinhardtii* isolated centrioles. Note the presence of procentrioles highlighted with the black arrowheads, **(b)** *d*STORM image of an isolated centriole. Scale bar: 250nm. **(c)** Confocal image of an expanded centriole using UltraExM (0.7%FA+1%AA). (d) Deconvoluted image in (c) using Hyvolution. (e-g) Plot profile of the regions indicated with a dotted white lane with the corresponding full width at half maximum (FWHM): *d*STORM≈57nm, UltraExM (confocal)~59nm, UltraExM (HyVolution)~21nm. (h-j) Plot profile from the polar transformation of the corresponding procentrioles quantified in (e-g). Scale bar in (c, d): 1μm.

Based on these results, we set out to develop a new method of expansion microscopy amenable to reveal fine architectural details while preserving the overall architecture of isolated organelles. We capitalized on the MAP protocol, which relies on a post-expansion staining of an intact proteome, thus providing a valuable starting point to optimize conditions to achieve our goal. We reasoned that a concentration of 4% PFA and 30% AA used in the original MAP protocol is too high for isolated centrioles and might induce inter-molecular crosslinking impeding expansion. Thus, we incubated centrioles in 1% FA/30% AA solution, but this resulted in centrioles that were neither fully expanded nor properly shaped (**Supplementary Fig. 1b**). We next omitted FA altogether from the solution and incubated centrioles in only 30% AA (**Supplementary Fig. 1b**). Under this condition, centrioles could not properly expand and appeared even more deformed preventing proper quantification of their diameter, indicating that FA treatment is a crucial step for linking amide groups of proteins to the acrylic polymer and cannot be omitted. We then tested whether the FA/AA step was necessary at all for centriole expansion. Thus, we incubated centrioles in PBS and observed that they could expand, but not as much as the polymer expansion factor and their shape looked severely deformed (**Supplementary Fig. 1b**). In contrast, we found that incubation in low percentage of FA alone, ranging between 0.3% and 1% FA resulted in expanded centrioles that appeared properly shaped (**Supplementary Fig. 1b and 1c**). In a second step, we tested whether lower concentrations of AA in combination with 0.3%-1% FA further improve the expansion performance (**Supplementary Fig. 1d**). We found that addition of 0.15% - 1% AA significantly improves the expansion of centrioles while 5% - 10% AA had no or little impact (**Supplementary Fig. 1d**). Our results demonstrate that a low formaldehyde (0.3-1% FA) and acrylamide (AA) concentration (0.15-1% AA) results in intact centriolar expansion with correct diameter. Importantly, this result indicates that this novel method that we termed <u>Ultra</u>structure <u>Exp</u>ansion <u>Mi</u>croscopy (UltraExM) preserves the ultrastructure of centrioles (Fig. 1f, 1j). Remarkably, UltraExM applied on isolated centrioles reveals unambiguously the nine-fold symmetry of the centriole as seen both for the α-tubulin and PolyE signal with correct diameters of 195 nm ± 2 nm and 225 nm ± 3 nm, respectively (Fig. 1f, 1i, 1j). In addition, lateral views of the UltraExM centrioles show a complete preservation of the centriolar shape as compared to cryoEM images of isolated centrioles (**Supplementary Fig. 2c-d**) and a perfect isotropic expansion of centrioles as demonstrated by the intact length-to-diameter ratio compared to non-expanded control samples (Fig. 1k). Moreover, we could reveal the central core decoration of the PolyE signal while retaining a complete tubulin decoration of the centriolar wall, indicating that we alleviated the antibody competition observed when using other expansion microscopy methods. Finally, we measured the roundness of centrioles, with a value of 1 reflecting a perfect round shape to monitor conceivable deformations of centrioles upon expansion. Here again, we found that the centriolar shape is best preserved in UltraExM compared to other expansion microscopy protocols (Fig. 1l). These findings indicate that UltraExM enables isotropic molecular expansion allowing the visualization of intact isolated macromolecular complexes.

### UltraExM surpasses *d*STORM

We then sought to test the potential of UltraExM by comparing confocal images of expanded centrioles imaged with a diffraction-limited microscope and non-expanded centrioles imaged with a super-resolution microscope. Therefore, we compared confocal images of expanded centrioles and non-expanded centrioles imaged by *d*STORM, both stained for PolyE. In order to accurately quantify the resolution in both modalities, we decided to image smaller structures such as procentrioles, which are assemblies with a height of 50 nm close to the mature centrioles^21^. As shown in Fig. 2a, UltraExM allows to clearly observe the procentrioles as fine structures that can be readily recognized and imaged on the side of mature centrioles. We next set out to image such centriole pairs by *d*STORM. Remarkably, *d*STORM imaging reveals mature centrioles and procentrioles as well as their associated connecting fibers (Fig. 2b and **Supplementary Fig. 3**). However, the 9-fold symmetric microtubule blades could not be visualized unambiguously (Fig. 2h). Strikingly, confocal images of UltraExM expanded samples reveal the 9-fold symmetrical pattern of procentrioles (Fig. 2c, 2i). These results indicate a higher labeling efficiency in UltraExM compared to *d*STORM experiments, probably due to limited antibody accessibility in the unexpanded state (Fig. 2c, f, i). Upon deconvolution of the images, the resolution can be further improved enabling for the first time the unequivocal visualization of the 9-fold symmetry of procentrioles with high precision (Fig. 2d, g, j). Overall, UltraExM combined with confocal fluorescence microscopy with deconvolution exhibits a high labeling efficiency and spatial resolution (Fig. 2e-g) enabling the characterization of ultrastructural components of macromolecular assemblies.

### Sub-microtubule triplet localization resolution of tubulin glutamylation

Polyglutamylation is a tubulin posttranslational modification that is critical for longterm stability of centriolar microtubules^22^. However the lack of a precise localization of this modification along the centriole prevents understanding how it could protect microtubule triplets. Therefore, we set out to analyze precisely its localization along the centriole, especially on the microtubule triplets using UltraExM (Fig. 3). Remarkably, UltraExM reveals for the first time that PolyE covers the outer surface of the tubulin signal, suggesting that it forms a shell around the centriole (Fig. 3a, arrowhead and Supplementary Videos 1 and 2). In addition we noticed a variation of PolyE decoration along the centriole with nine discrete puncta at both proximal and distal ends of the central core region (Fig. 3a). To gain more insight into PolyE distribution and to prevent any artifact due to the anisotropic resolution of confocal microscopy, we next performed an isotropic 3D reconstruction (Fig. 3b-f)^23^. By combining over fourteen mature centrioles, we generated an isotropic averaged volume of the centriole (Fig. 3b-f). This result confirmed that PolyE fully covers the outer surface of the central core region of mature centrioles (Fig. 3e), while it depicts nine discrete signals at both distal (Fig. 3d) and proximal (Fig. 3f) ends of this region. By measuring the diameters of both PolyE and α-tubulin signals, we found that PolyE depicts a measured diameter of 88-140 nm larger than that of tubulin (**Supplementary Fig. 4a-d**). Our observation suggests that tubulin polyglutamylation shelters the outer surface of centriolar microtubules thus potentially stabilizing centrioles.

**Figure 3.**
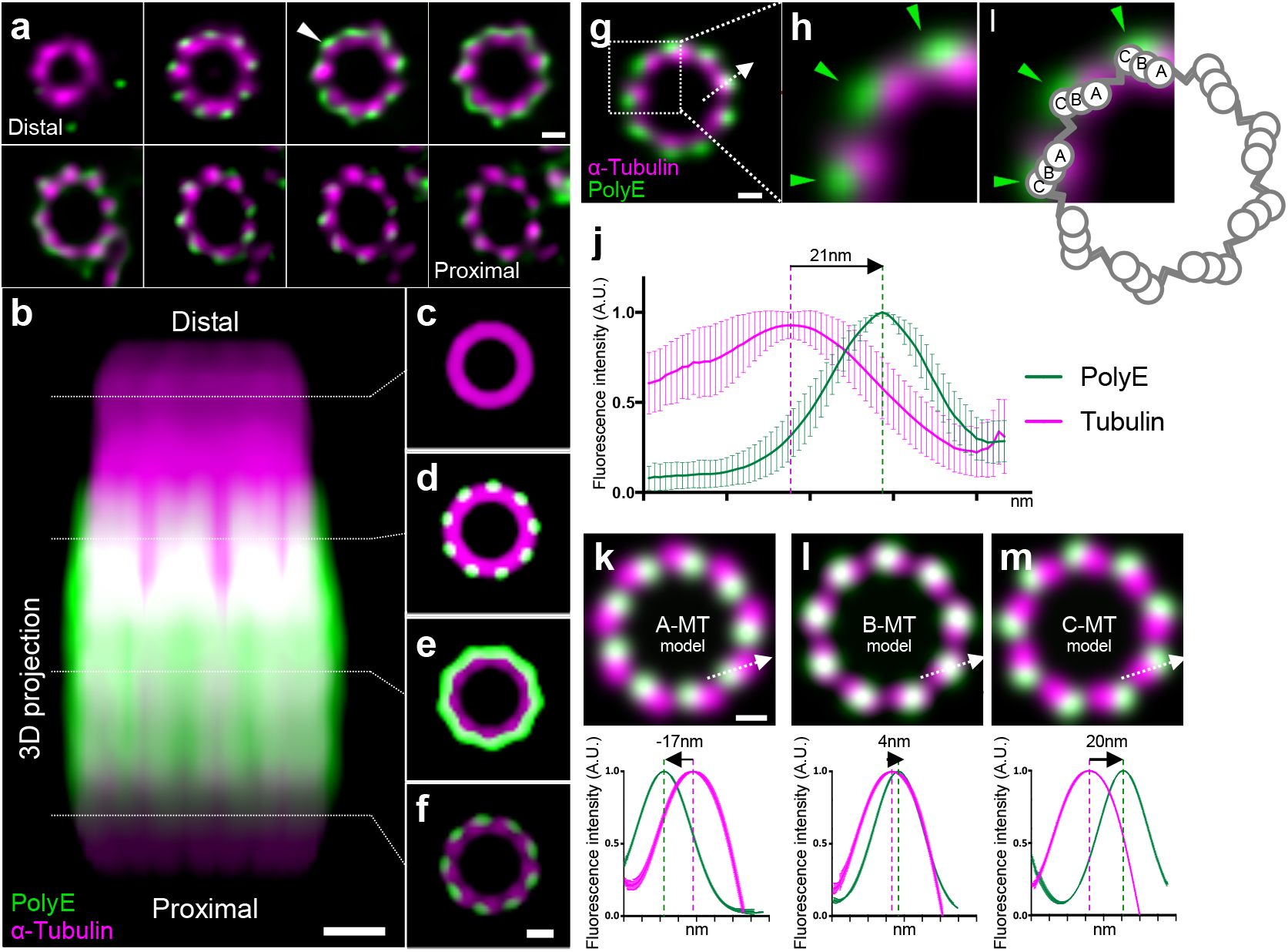
Sub-triplet localization revealed by UltraExM. **(a)** Gallery of a Z stack from distal to proximal of a mature centriole expanded using UltraExM (0.7%FA+1%AA) and stained with α-tubulin (magenta, Alexa568) and PolyE (green, Alexa488). Scale bar: 200nm. Arrowhead points to PolyE surrounding the tubulin signal, **(b)** 3D volume reconstruction of a mature *Chlamydomonas* centriole stained with α-tubulin (magenta, Alexa568) and PolyE (green, Alexa488). n=7. Scale bar: 200nm. **(c-f)** Sections through the centriole spanning the proximal **(c)** and the central core **(d-f)** regions. Scale bar: 300 nm. **(g-i)** Representative image taken from a Z-stack of the distal most-part of the central core of a deconvolved mature centriole stained for α-tubulin (magenta, Alexa568) and PolyE (green, Alexa488). Scale bar: 200nm. Green arrowheads highlight the PolyE signal **(h, i).** The dotted arrow illustrates how the fluorescence intensity was measured in **j. (i)** Schematic representation of microtubule triplets superimposed onto the fluorescent signal shown in H. (j) Quantification of the fluorescence intensity shift between the magenta and the green signal. Note that the shift is of 21 nm (n=61). (k-m) Model representation of PolyE signal onto specific microtubule blades, A- (k), B- (l) and C- (m). Note that in this model, the entire triplet is stained with α-tubulin (magenta). Below is represented the fluorescence peak shift between the two colors in each condition. Scale bar: 200nm.

Next, we investigated the precise localization of the nine discrete puncta of PolyE signal at the distal region. Interestingly, it has been shown by electron microscopy that the cilium, composed by only A- and B- microtubules, is polyglutamylated solely on the B-microtubule^24^. We then asked whether these PolyE puncta could correspond to a similar sub-microtubule triplet localization. Therefore, we focused on the most-distal part of the central core region of individual mature centrioles treated with UltraExM (Fig. 3g). Remarkably, we found that the polyglutamylated signal did not co-localize with the tubulin signal but instead showed a marked shift compared to the center signal of tubulin (Fig. 3g-i). We measured the fluorescence signal shift between the tubulin and PolyE signals and found that the PolyE signal is displaced by 21 nm from the tubulin signal (Fig. 3j). To identify to which of the A-, B- or C-microtubule triplets PolyE could be assigned, we modeled its position on each microtubule (Fig. 3k-m and **Supplementary Fig. 4e-m**) and measured the displacement of the expected PolyE from the tubulin signal. Strikingly, the results demonstrate that deposition of PolyE on the C microtubule gives a displacement of ~ 20 nm, similar to the one observed in the UltraExM experiments (Fig. 3j). This establishes that UltraExM is able to reveal exquisite localization pattern and to distinguish a C-microtubule triplet localization for polyglutamylated tubulin in mature centrioles.

**Figure 4.**
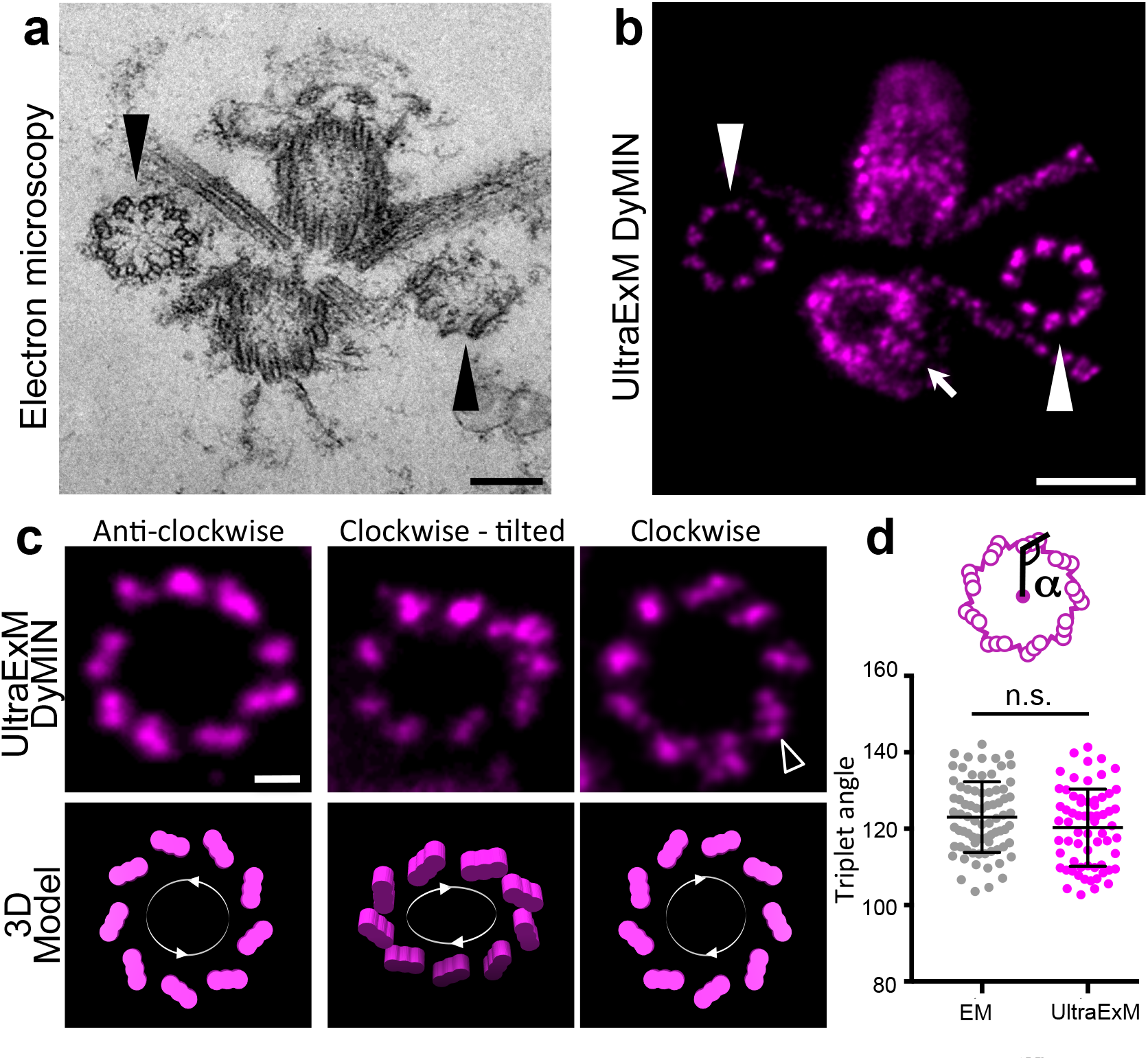
UltraExM combined to STED reveals the centriole chirality. **(a)** Electron microscopy (EM) image of a centriole pair comprising two mature centrioles and two procentrioles (black arrowheads) interconnected by striated fibers. Scale bar: 200nm **(b)** A similar centriole pair stained for α-tubulin (magenta, STAR RED) after UltraExM (1% FA) and imaged using DyMIN. Note that the overall ultrastructure of this organelle resembles the EM image. White arrowheads points to procentrioles and the arrow points to the mature centriole. Scale bar: 1 μm **(c)** Representative DyMIN images of procentrioles stained for α-tubulin (magenta, STAR RED) highlighting their anticlockwise or clockwise orientations. Below is the interpretation of such orientations in a 3D schematic model. Black arrowheads points to individual blades within a microtubule triplet. Scale bar: 200nm **(d)** Quantification of the angle between the center of the centriole and the microtubule triplet both from EM (123° ± 1°) and DyMIN (120° ± 1°) images. P=0.0912

### Unveiling centriole chirality by combining UltraExM with STED-microscopy

To further investigate the ability of UltraExM to reveal the molecular architecture of centrioles, we combined UltraExM with STED microscopy either single (Fig. 4) or dual color (**Supplementary Fig. 5 and Supplementary Video 3**). A key prediction of the macromolecular expansion of the centriole is that the microtubule triplet chirality, an evolutionary conserved feature of the centriole, should now become apparent using tubulin as a marker. As shown by electron microscopy, centrioles are composed of nine microtubule triplets with a characteristic angle and are arranged in a clockwise manner as seen from the proximal side (Fig. 4a). UltraExM-treated centriole pairs were imaged using DyMIN, a STED-microscopy imaging method relying on an intelligent light dose management^25^. Remarkably, the resolution improvement allows a glimpse of the triplet structure of microtubules on the procentrioles (Fig. 4b-c, arrowheads and **Supplementary Fig. 5**). In addition, the images give also an insight into microtubule triplets of the mature centrioles (Fig. 4b, arrow). Strikingly, UltraExM-STED enables also visualization of the clockwise and anticlockwise orientation of microtubule triplets in procentrioles, respectively (Fig. 3c). In addition, we could even occasionally identify three distinct fluorescent peaks for some microtubule triplets that possibly correspond to the A-, B- and C-microtubule, respectively (Fig. 4c, arrowhead). Furthermore, we measured the angle between the center of the procentriole and the microtubule triplet and compared it to the angles measured by electron microscopy (Fig. 4d). We found a similar angle of ~120° corroborating that STED-microscopy can visualize the triplet orientation of UltraExM-expanded centrioles and indicating that UltraExM preserves the nanometric conformation of the sample.

Together, these results demonstrate that UltraExM combined with super-resolution microscopy methods can reveal ultrastructural features with unprecedented detail such as the microtubule triplet chirality that was only accessible by electron microscopy until now.

## DISCUSSION

In summary, we report a novel method of expansion microscopy, UltraExM, amenable to reveal fine ultrastructural details of multiprotein complexes as exemplified with isolated centrioles. UltraExM reveals unambiguously the 9-fold symmetry of centrioles, both for tubulin and polyglutamylated tubulin signals using confocal microscopes. Moreover, we found that coupling UltraExM with STED imaging unveils for the first time the chirality of the microtubule triplets as well as distinguishes between the A, B and C microtubules within the blades.

The ability of super-resolution microscopy to resolve the molecular architecture of multiprotein complexes or macromolecular assemblies is controlled by several factors such as fixation, protein labeling density and size of the fluorescent probes. UltraExM shows new paths to overcome these limitations by using near-native expansion of the macromolecule prior to labeling. Using only low concentrations of formaldehyde and acrylamide for fixation, macromolecules are only minimally structurally affected. In addition, the post-labeling on the expanded sample improves epitope accessibility for immunolabeling and thus the labeling density. Moreover, as the overall size of the expanded macromolecule is ~4 -fold larger, labeling results in a likewise smaller linkage error. Thus, even larger fluorescent probes such as primary and secondary IgG antibodies can be used advantageously to reveal molecular details of macromolecular assemblies.

Expansion microscopy methods covalently anchor fluorescent probes or proteins directly into a polymer network that isotropically expands through dialysis in water^15–18^and thus can overcome several limitations of super-resolution microscopy and enable imaging of fixed cells and tissues with a spatial resolution of ~70 nm^15^. Despite these advances, the reliability of the isotropic expansion process at the molecular level remained elusive. Our results show unambiguously that refined ExM protocols such as UltraExM preserve ultrastructural details and can thus be used successfully to visualize the molecular architecture of multiprotein complexes.

Combined with super-resolution microscopy methods as illustrated here for STED microscopy, UltraExM can reveal unprecedented structural features that were so far only accessible by electron microscopy. Of course, there is ample of scope for further improvements but our data show unambiguously that UltraExM has what it takes to occupy an important place among the methods useable for structure elucidation of macromolecular assemblies.

## METHODS

Methods, including statements of data availability and any associated accession codes and references, are available in the online version of the paper.

## ACKNOWLEDGMENTS

We thank Nikolai Klena for critical reading of the manuscript. DG and M.C are supported by the European Reaserch Council ERC StG 715289 and PG and VH by the Swiss National Science Foundation (SNSF) PP00P3_157517. FZ and MS acknowledge support by the Deutsche Forschungsgemeinschaft (DFG) within the Collaborative Research Center 166 ReceptorLight (projects A04 and B04). M.U. is supported by the ERC (GA No 692726 GlobalBioIm).

## AUTHOR CONTRIBUTIONS

D.G, F.U.Z, M.S, V.H and P.G. conceived and designed the project. M.S, V.H and P.G supervised the project. D.G and F.U.Z performed all ExM experiments. DG performed all UltraExM experiments with the help of S.B. and the data analysis. F.U.Z. performed the *d*STORM imaging. J.H., G. S. and M.R. performed the STED imaging. D.F. and M.U. performed the 3D averaging. M. C. initiated the UltraExM project. E.B. helped in setting up ExM. All authors wrote and revised the final manuscript.

